# Identifying and Addressing Systematic Data Leakage in Protein-Ligand Affinity Benchmarks

**DOI:** 10.64898/2026.06.29.735309

**Authors:** Björn Mattsson, W. Patrick Walters

## Abstract

Accurate prediction of protein-ligand binding affinity is a crucial goal in structure-based drug discovery, with the potential to significantly shorten development timelines. Recently, a new wave of machine learning models based on co-folding, such as Boltz-2 and IsoDDE, has demonstrated performance that matches or exceeds that of gold-standard physics-based methods like Free Energy Perturbation (FEP). This paper provides a critical assessment of these claims, revealing that current benchmarks are heavily influenced by data leakage, and proposes a new benchmark that explicitly controls for data leakage.

We demonstrate that splitting by protein-sequence identity is inherently insufficient to prevent data leakage due to “target mirroring,” in which homologous proteins with low overall sequence identity still exhibit highly correlated binding profiles. Our meta-analysis of documents in the ChEMBL 36 database identifies more than 6,000 such assay pairs and finds that leakage persists for sequence-identity thresholds as low as 0.2, well below the values commonly used in benchmarks today. Additionally, we show that a ligand-only baseline model, which lacks protein structural information, achieves surprisingly high performance on the FEP+ 4 and OpenFE benchmarks (*r* = 0.66 and *r* = 0.36, respectively).

Our results indicate that current benchmarks tend to reward models for memorizing training data and exploiting localized leakage rather than truly learning biophysical principles. To address this issue, we propose the Novelty-Tiered Affinity Benchmark, in which the test data is partitioned into ligand novelty tiers. In the most challenging tier (Tanimoto similarity *<* 0.35), ligand-only models perform notably worse (*r* = 0.14), offering a clear baseline for evaluating genuine generalization. We argue that the field must move beyond sequence-based splits to ensure that AI-driven discovery translates into successful prospective laboratory research.

## 1 Introduction

Robust machine learning (ML) requires a rigorous separation between training and evaluation data to ensure that a model is truly learning underlying principles rather than simply memorizing inputs. When test data overlaps significantly with the training set, a phenomenon known as data leakage, performance metrics become artificially inflated [1]. In such cases, high accuracy reflects the model’s ability to interpolate within known data rather than its capacity to generalize to novel chemical space.

This issue is a persistent challenge across the broader field of machine learning in drug discovery. For example, in QSAR (Quantitative Structure-Activity Relationship) modeling, random splits [2] often yield overly optimistic results because models can readily exploit similarities between closely related compounds in the training and test sets. Similarly, in docking benchmarks such as PDBBind [3], the inclusion of homologous structures can allow models to “cheat” by referencing known solutions rather than uncovering genuine molecular interactions [4, 5].

Despite these known pitfalls, accurately predicting protein-ligand binding affinity remains a central “Holy Grail” in structure-based drug discovery. The ability to predict how strongly a small molecule binds to a target protein allows researchers to prioritize which compounds to synthesize and test. This can significantly compress drug discovery timelines and reduce the prohibitive costs associated with early-stage discovery. The potential value of this technology has attracted substantial interest from machine learning researchers, both within academia and industry, leading to the emergence of several datasets as de facto benchmarks.

However, many of these “gold-standard” benchmarks have recently attracted considerable criticism for failing to prevent the very leakage described above. DUD-E [6], for several years one of the most widely used benchmarks for docking-based virtual screens, has been shown to exhibit substantial leakage when used with machine learning models [7, 8, 9, 10]. CASF-2016 [11], another prominent benchmark, was originally developed to evaluate physics-based scoring functions with few parameters. When modern machine learning models with millions of parameters began using CASF-2016, typically while training on PDBBind [12], this intrinsic shortcoming surfaced as inflated results because nearly identical data points appeared on both sides of the train/test split [10, 13]. These failures suggest that, as a field, we have not adopted robust benchmark strategies in which high performance cannot be achieved by memorization. Consequently, it is often not possible to assess, from published results alone, whether models would be useful in prospective drug discovery research.

In this paper, we critically analyze benchmarks based on protein-identity splits, which have recently been used to claim that ML methods surpass physics-based methods, and we contribute a more robust benchmarking suite. The paper starts with an overview of recent claims by co-folding methods in Section 2. In Sections 3 and 4, we present a case study and a meta-analysis that explain why protein-sequence-identity-based splits lead to data leakage. Sections 5 and 6 contain concrete evidence of data leakage problems in the FEP+ 4 and OpenFE benchmarks. In Section 7, we present a new benchmark methodology for the field, inspired by Runs N’ Poses [14]. Section 8 presents our conclusions, and Section 9 links to our benchmark dataset and the code.

This criticism targets flawed benchmarks and the claims made with them, not the models themselves. To better understand the performance characteristics of different binding affinity prediction methods and the chemical space in which they are useful, the field needs robust benchmarking methods that control for data leakage.

## 2 Co-folding and Sequence Similarity Cutoffs

In recent months, the scientific community has seen significant interest surrounding the ability of co-folding methods to predict binding affinity. While initial results from models like Boltz-2 [15], IsoDDE [16], and AQAffinity [17] appeared promising, a closer look reveals a fundamental flaw. The accompanying headlines and abstract reported results using a data-splitting strategy with a 90% sequence similarity cutoff, a threshold that leads to substantial data leakage. As a result, what appears to be advanced “prediction” could, on parts of these benchmarks, be achieved by little more than a simple similarity search (see Sections 5 and 6). It should be noted that both Boltz-2 and AQAffinity presented more nuanced results as well: Boltz-2 reported challenges reaching as promising results on internal Recursion targets, and AQAffinity reported two assays filtered to 70% sequence similarity and 70% ligand similarity, where neither the AQAffinity nor the Boltz-2 method was able to rank the compounds by binding affinity.

The IsoDDE paper also validated its performance on a separate benchmark constructed from newly deposited assays in ChEMBL 35. The use of recently deposited data is not in itself sufficient; newer assays often report on compounds and targets that were already extensively characterized in earlier versions of the database. The authors provided a plot suggesting equivalent performance across “high,” “medium,” and “low” similarity bins. However, this analysis on a different dataset does not preclude the possibility that the model exploited data leakage to achieve the inflated results on which its central conclusions were based.

The challenge in activity prediction benchmarks lies in the correlation between related targets. For protein families such as kinases or GPCRs, a ligand often exhibits similar binding profiles across multiple homologs. If a dataset includes such related proteins, a model can achieve high accuracy by essentially retrieving known activity values from a training homolog to “predict” the activity for a test target. This data mirroring undermines the evaluation of a model’s true predictive power.

Several efforts have been made to address these limitations through more sophisticated splitting strategies. PDBBind CleanSplit [5] uses a combination of protein similarity (TM-score), ligand similarity (Tanimoto similarity), and pocket-aligned RMSD to ensure independence. Similarly, PLINDER [18] incorporates overall protein similarity, binding site similarity, and ligand similarity into its benchmark design. Leakproof PDBBind [4] rigorously cleans the data before splitting it based on sequence similarity, ligand chemical similarity, and structural interaction patterns. The primary disadvantage of these structure-aware approaches is their reliance on available crystallographic or docked data. In many practical scenarios, researchers seek to train on large-scale databases like ChEMBL or BindingDB, where 3D structural information or exact binding modes are often unknown. While there are alternative approaches that do not require structural information, their splitting criteria often prove insufficient for preventing the various forms of leakage discussed herein.

It is worth noting that the activity prediction methods used in these modern co-folding approaches bear a striking resemblance to proteochemometric (PCM) methods [19], which have been popular for the last two decades. In PCM approaches, a combined representation of the protein and ligand is used to train a model that predicts the activity of new compounds across multiple targets simultaneously. However, the pitfalls of this paradigm were highlighted in a 2025 study by the Isayev group [20], which identified numerous data leakage issues common to these models. The authors found that permuting the protein representations often had minimal impact on model performance, suggesting that predictions were largely driven by ligand representations and preexisting statistical correlations rather than by an understanding of specific protein-ligand interactions.

## 3 Case Study: Correlation Among Muscarinic Receptors

To examine the real-world implications of “target mirroring,” we analyze a series of compounds tested across the five human muscarinic acetylcholine receptors: M1, M2, M3, M4, and M5. This data was extracted from a single 1998 paper [21] in which the authors were optimizing a chemical series of antagonists specifically to be selective for the M4 receptor over the M1 receptor. In medicinal chemistry, achieving such selectivity among these subtypes is paramount; for example, while M4 antagonism can help restore dopamine-acetylcholine balance in Parkinson’s disease, blocking the closely related M1 receptor can lead to serious side effects, including cognitive fog and memory loss.

The ChEMBL database reports IC_50_ values for the compounds from this optimization campaign across the five subtypes. Our analysis, visualized in Figure 1, reveals that in 5 out of 10 pairwise comparisons, the Pearson *r* is *≥* 0.77 (equivalent to an *R*^2^ *≥* 0.6). Specifically, correlations reach as high as *r* = 0.90 for M1 vs. M3 and *r* = 0.96 for M1 vs. M5. This indicates that if a model is trained on one muscarinic subtype, it can often simply “look up” the activity for another subtype based on the established correlation observed within the chemical series.

**Figure 1:**
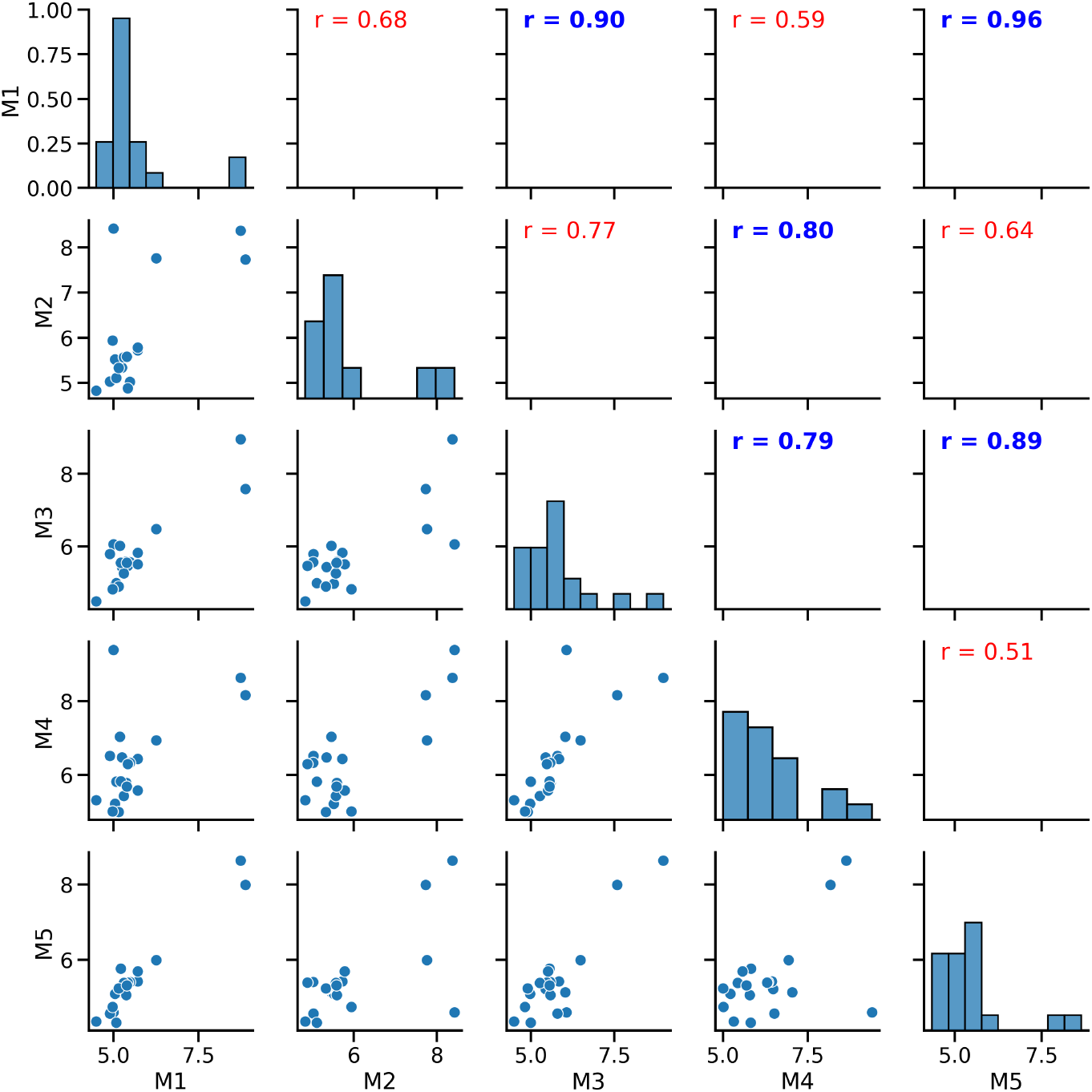
pIC_50_ correlation matrix of five muscarinic assays. Data sourced from Augelli-Szafran (1998) [21]. The upper triangle displays Pearson correlation coefficients (*r*), with values *≥* 0.77 highlighted in blue and values *<* 0.77 in red.

Crucially, the overall sequence similarity between any two muscarinic receptors typically ranges between 40% and 65%. These values are significantly lower than the 90% similarity cutoff often used to partition training and test sets in affinity prediction benchmarks. Under such benchmarks, the model would encounter these highly correlated targets across the split boundary, enabling high “predictive” scores from simple data retrieval of training homologs rather than an understanding of M4-specific selectivity.

## 4 Meta-Analysis of Assay Correlation in ChEMBL

To explore how common this data leakage is, we reviewed all documents in the ChEMBL 36 database [22] to identify pairs of assays from the same paper meeting four criteria:

1. **Multiple Assays:** A paper reported data from at least two distinct assays.
2. **Compound Overlap:** At least 10 compounds tested in both assays with reported pChEMBL values.
3. **Dissimilarity:** Less than 90% overall sequence similarity between assay targets.
4. **Statistical Correlation:** An *R*^2^ of at least 0.6 between assay values.

Table 1 provides details of this analysis. Note that in 99% of cases, the overall sequence similarity between assay targets was *<* 0.9. We found 6,115 assay pairs that meet these criteria, covering a wide variety of targets, including kinases, GPCRs, and metalloenzymes. A sample of the correlated assays is shown in Table 2. The full table, as well as the scripts used to perform the analysis, are available in the supporting materials for this paper. This type of study is standard in medicinal chemistry; teams spend significant time seeking selectivity for homologs with similar binding pockets. When benchmarks ignore these correlations, they inadvertently reward models for memorization.

**Table 1:** An analysis of the number of assays in ChEMBL 36 with correlated values.

| Criteria | Assay Pairs | Papers |
| --- | --- | --- |
| At least 2 assays with at least 10 compounds in common | 59,552 | 8,454 |
| Sequence similarity $< 0.9$ | 59,085 | 8,218 |
| $R^2 \geq 0.6$ | 6,115 | 2,689 |

**Table 2:** A sample of the 6,115 assay pairs with correlated values found in the ChEMBL 36 database.

| Target 1 | Target 2 | pChEMBL $R^2$ | Sequence Similarity |
| --- | --- | --- | --- |
| Calcium/calmodulin-dependent protein kinase type II subunit gamma | Serine/threonine-protein kinase 3 | 0.73 | 0.20 |
| Histone deacetylase 1 | Histone deacetylase 4 | 0.90 | 0.11 |
| G-protein coupled receptor 183 | G-protein coupled receptor 35 | 0.69 | 0.24 |
| Serine/threonine-protein kinase 10 | Calcium/calmodulin-dependent protein kinase type 1G | 0.63 | 0.17 |
| Acyl-CoA desaturase 1 | Stearoyl-CoA desaturase | 0.82 | 0.85 |
| Muscarinic acetylcholine receptor M3 | Muscarinic acetylcholine receptor M5 | 0.74 | 0.53 |
| Collagenase 3 | 72 kDa type IV collagenase | 0.80 | 0.41 |
| Carbonic anhydrase 1 | Carbonic anhydrase 4 | 0.70 | 0.27 |
| Carbonic anhydrase 3 | Carbonic anhydrase 5A, mitochondrial | 0.89 | 0.44 |
| 3',5'-cyclic-AMP phosphodiesterase 4B | 3',5'-cyclic-AMP phosphodiesterase 4C | 0.71 | 0.64 |
| 5-hydroxytryptamine receptor 2C | 5-hydroxytryptamine receptor 2B | 0.67 | 0.43 |
| Muscarinic acetylcholine receptor M2 | Muscarinic acetylcholine receptor M4 | 0.82 | 0.59 |
| Acetylcholinesterase | Acetylcholinesterase | 0.92 | 0.59 |
| Hepatocyte growth factor activator | Suppressor of tumorigenicity 14 protein | 0.77 | 0.22 |
| Muscarinic acetylcholine receptor M2 | Muscarinic acetylcholine receptor M1 | 0.84 | 0.45 |
| MAP kinase-interacting serine/threonine-protein kinase 2 | NUAK family SNF1-like kinase 2 | 0.64 | 0.20 |
| Prolyl hydroxylase EGLN2 | Egl nine homolog 1 | 0.83 | 0.46 |
| Sphingosine 1-phosphate receptor 1 | Sphingosine 1-phosphate receptor 2 | 0.85 | 0.47 |
| Phosphorylase b kinase gamma catalytic chain, skeletal muscle/heart isoform | Interleukin-1 receptor-associated kinase 4 | 0.76 | 0.15 |
| Calcium/calmodulin-dependent protein kinase type II subunit delta | Serine/threonine-protein kinase 3 | 0.74 | 0.20 |

## 5 Deconstructing the FEP+ 4 and OpenFE Benchmarks

Many reports on affinity prediction using co-folding methods compare predictions with those from alchemical free-energy calculations. The Boltz-2 team introduced two benchmarks for this purpose: FEP+ 4 (with targets TYK2, CDK2, JNK1, p38*α*) and the OpenFE subset of the Ross et al. 2023 benchmark (with 39 targets) [15], using data originally designed to validate FEP methods [23, 24].

To prevent data leakage, the Boltz-2 team excluded targets with *≥* 90% sequence identity to any test target from the training set. Using this standard, the Boltz-2 team reported an average Pearson correlation coefficient (*r*) of 0.66 over the FEP+ 4 targets and 0.62 on the OpenFE targets, claiming to approach the performance of FEP methods (*r* = 0.78 and 0.72, respectively) [15]. The Isomorphic Labs team later reported average correlations of 0.85 and 0.73 on the same benchmarks, asserting that IsoDDE’s affinity predictions surpass those of gold-standard physics-based methods [16].

We reproduced the FEP+ 4 and OpenFE data split by clustering protein sequences at *≥* 90% (see Appendix A for details). This revealed that targets with nearly identical binding pockets can end up in different splits: for example, JNK1 and JNK2, which share 81% sequence identity. Consequently, the compound lig_18633-1 (CHEMBL426134) appears in both the test set (JNK1, pIC_50_ = 6.75) and the train set (JNK2, pIC_50_ = 6.56), with predicted binding poses that are virtually indistinguishable (see Figure 2, left).

**Figure 2:**
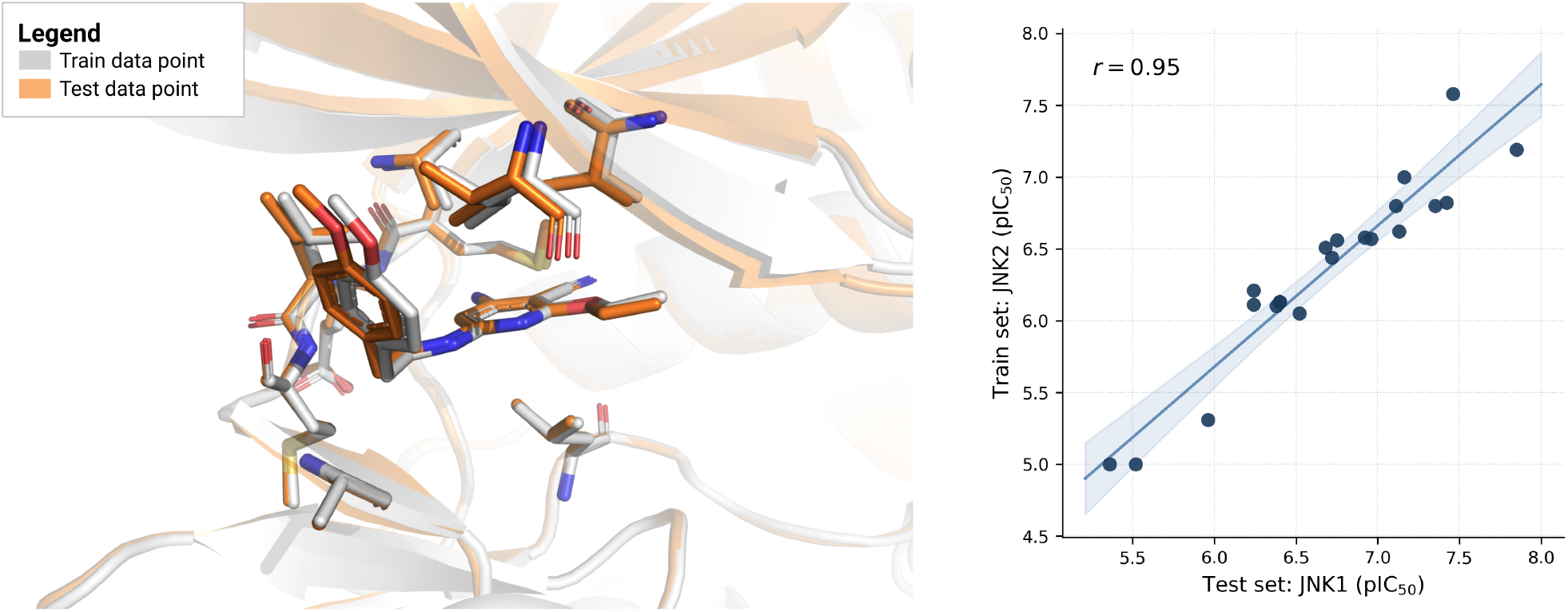
Data leakage in the FEP+ 4 benchmark. *Left:* Predicted binding poses of lig_18633-1 co-folded with JNK1 (test set, orange, pIC_50_ = 6.75) and JNK2 (train set, gray, pIC_50_ = 6.56) using Boltz-2, with residues within 3.5 Å shown. The poses are virtually indistinguishable, illustrating that JNK1 and JNK2 present the same binding environment for this compound series. *Right:* Pearson correlation (*r* = 0.95) between pIC_50_ values for the full JNK1 (test) and JNK2 (train) compound series. Because JNK1 and JNK2 share 81% sequence identity, the data split methodology places them in different splits despite their near-identical binding pockets.

We find that the entire JNK1 compound series (assay CHEMBL860181) has a corresponding JNK2 measurement in the training set (assay CHEMBL860184). Both series were published in the same 2006 paper [25], and the Pearson correlation between the JNK1 test series and the JNK2 train series is *r* = 0.95. A model that simply looks up JNK2 training values to “predict” JNK1 would achieve this correlation without any generalization (see Figure 2, right).

This issue is not exclusive to JNK1. Each FEP+ 4 test target has multiple closely related targets in the training split (Table 3). The JNK1/JNK2 case is the clearest example of leakage: the entire compound series appears on both sides of the split. For other test targets, the overlap is partial, with several compounds from the test series or their close analogs present in the training set, tested against targets with similar binding pockets.

**Table 3:** Targets with similar binding pockets to each FEP+ 4 test target, grouped by data split. For each test target, closely related targets are present in both the test and train sets, indicating that the protein sequence identity-based split leads to many opportunities for data leakage.

| Test target | In test set | In train set |
| --- | --- | --- |
| TYK2 | TYK2, JAK1, JAK2, ITK, BTK, EGFR, CDK | JAK3, SRC |
| CDK2 | CDK2, CDK8 | CDK1, CDK3, CDK4, CDK5, CDK6, CDK9, DYRK1A |
| JNK1 | JNK1, JNK3, p38α, CDK2 | JNK2, p38β, ERK2, ERK1, DYRK1A |
| p38α | p38α, JNK1, CDK2 | p38β, p38γ, p38δ, JNK2, ERK2, ERK1, DYRK1A |

We find similar data leakage problems in the OpenFE subset when using a 90% sequence identity threshold. The OpenFE set is divided into several subsets. Although we have investigated the entire collection, we will refer to two specific subsets, “merck” and “miscellaneous”. For the merck TNKS2 test data series, we find that 19 of 27 ligands appear in the CHEMBL4322251 assay against TNKS1, which shares only 71.4% sequence identity with TNKS2. This second assay appears in the training set even though the correlation between the two series is *r* = 0.98. Similarly, for the miscellaneous BTK test data series, we find all 6 compounds appearing in the assay CHEMBL3588238, which measures inhibition of LCK. While these targets only share 22.5% sequence identity and therefore end up in different data splits, their Pearson correlation is *r* = 0.94. See Figure 6 in Appendix B for the corresponding plots.

## 6 Ligand-only Baselines on the FEP+ 4 and OpenFE Benchmarks

To assess the impact of this leakage, we developed ligand-only machine learning baselines that focus exclusively on ligand structure and exclude a protein-binding component. These baselines used learned molecular representations from Chemprop [26], molecular fingerprints, and normalized molecular descriptors as inputs, with the protein target represented by a one-hot encoding that contains no protein structural information (see AppendixC for details on the ML method). The baselines were trained on ChEMBL 36 [22], which had been split following the FEP+ 4 and OpenFE benchmark methodology. For decades, ligand-based models have been used in the computational chemistry community to find analogs of known hits. The main drawback is that, because they are only implicitly aware of the binding domain, they are not designed to discover novel chemistry. Instead, in these scenarios, the field has relied on physics-based methods, which, under the right conditions, have identified novel binders.

On the FEP+ 4 test set, the best ligand-only model matched Boltz-2 and the physics-based OpenFE model at *r* = 0.66, and a model that uses just twelve molecular descriptors achieved *r* = 0.49 (see Figure 3). On the OpenFE benchmark, the best ligand-only model achieved *r* = 0.36, with a larger gap between the ligand-only models and the other structure-based machine learning models. The ligand-only methods achieve higher performance than one would expect in true de novo situations (*r <* 0.20; see the left-hand side of Figure 4). This shows that any ML method exploiting this leakage would achieve inflated results, invalidating comparisons with physics-based methods that operate without training data. Furthermore, more diverse training data would likely lead to greater opportunities for data leakage when using protein-sequence-based data splitting. Boltz-2, for example, in addition to ChEMBL relies on BindingDB [27] and three other data sources.

**Figure 3:**
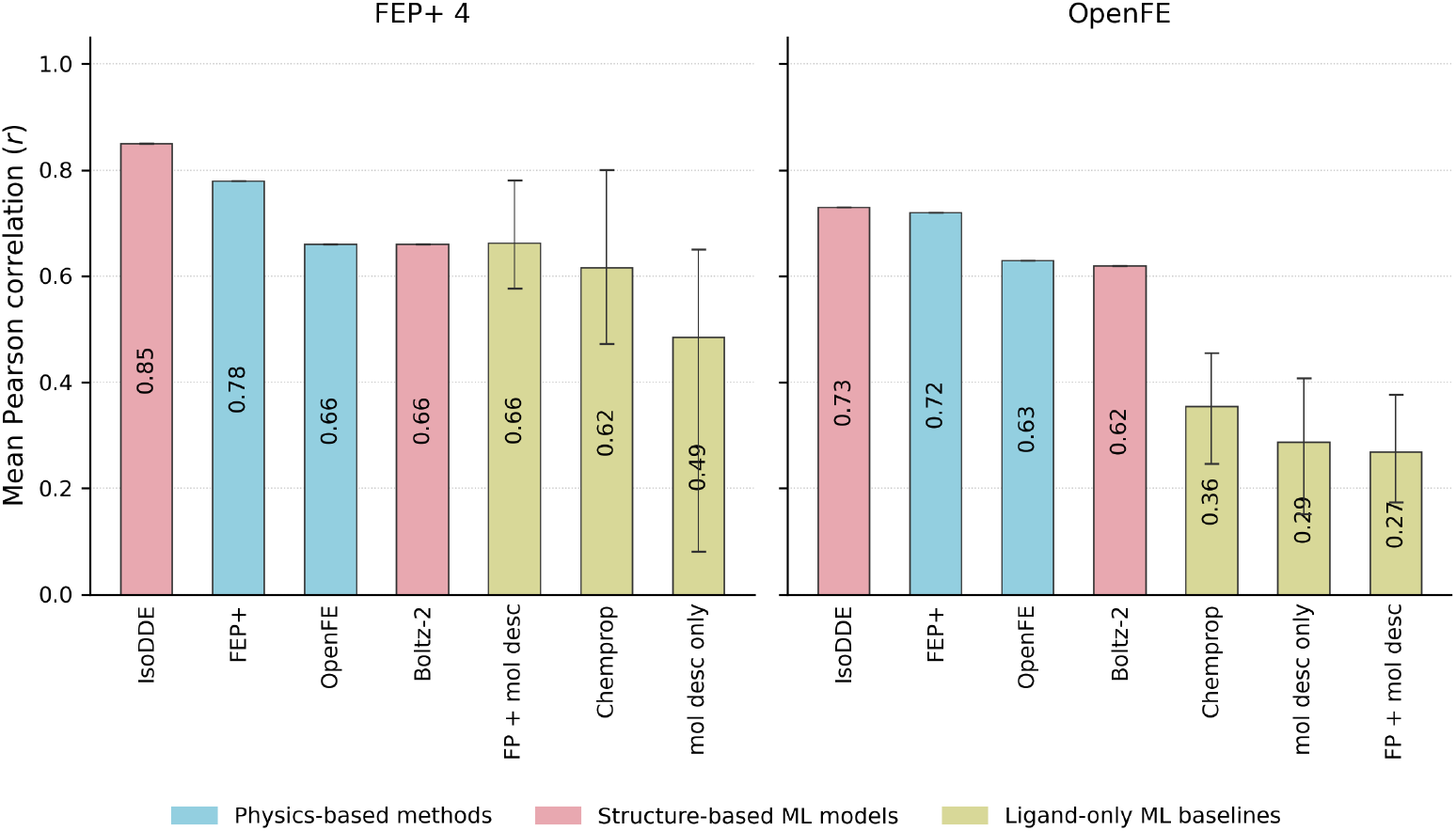
Mean Pearson correlation (*r*) on the FEP+ 4 and OpenFE benchmarks for published methods and ligand-only baselines. The mean was calculated as a size-weighted average to align with previously reported results. The error bars show bootstrapped 95% confidence intervals for the ligand-only methods. Results for methods other than the baselines were taken from the IsoDDE report [16].

**Figure 4:**
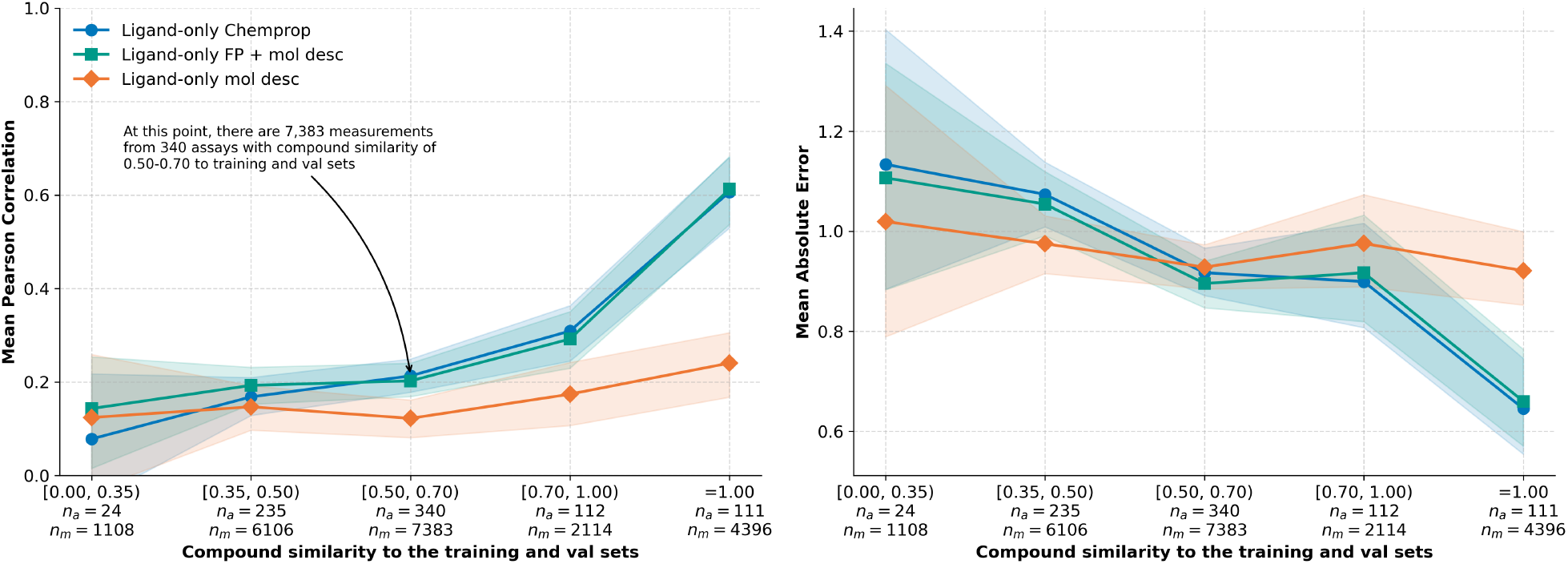
Performance of ligand-only baseline models for different ligand novelty tiers. Higher similarity to training compounds indicates greater data leakage, reflected in higher ligand-only model performance. Each bin along the x-axis groups compounds by their maximum Morgan Fingerprint (2048-bit, radius 2) Tanimoto similarity to training or validation compounds. Compounds with an exact match to a training or validation compound are separated out into their own bin (=1.00). *n*_*a*_ is the number of assays and *n*_*m*_ the number of measurements within that bin. The mean was calculated as an unweighted average across all assays in that bin, and shaded areas show bootstrapped 95% confidence intervals.

Even though we conclude that the performance of machine learning methods can be significantly inflated on this benchmark by exploiting the substantial data leakage, the evidence presented here is insufficient to conclude that Boltz-2 and IsoDDE derive the majority of their performance from memorizing ligand structures. To separate the contribution from ligand memorization and genuine generalization, we need better benchmarks, which the next section will focus on.

## 7 Proposed Splitting with Ligand Novelty

Unless one is extremely careful, any machine learning model trained to predict binding affinity is highly vulnerable to data leakage. Merely adjusting the overall sequence similarity threshold is insufficient. In many cases, we encounter proteins with high binding-site similarity but low overall similarity. The team that created Leakproof PDBBind recommends a sequence similarity cutoff of 0.5 and a ligand similarity cutoff of 0.99 [4]. To explore the connection between overall sequence similarity and data leakage, we varied the similarity cutoff in the calculations in Section 4 from 0.1 to 0.9. The plot in Figure 5 displays the results, with overall sequence similarity on the x-axis and the number of assay pairs showing correlated activity on the y-axis. Even with a sequence similarity cutoff of 0.5, we find more than 4,000 assay pairs reporting correlated activity across different targets. This result, along with a similarity cutoff that captures only identical molecules, calls the Leakproof PDBBind criteria into question.

**Figure 5:**
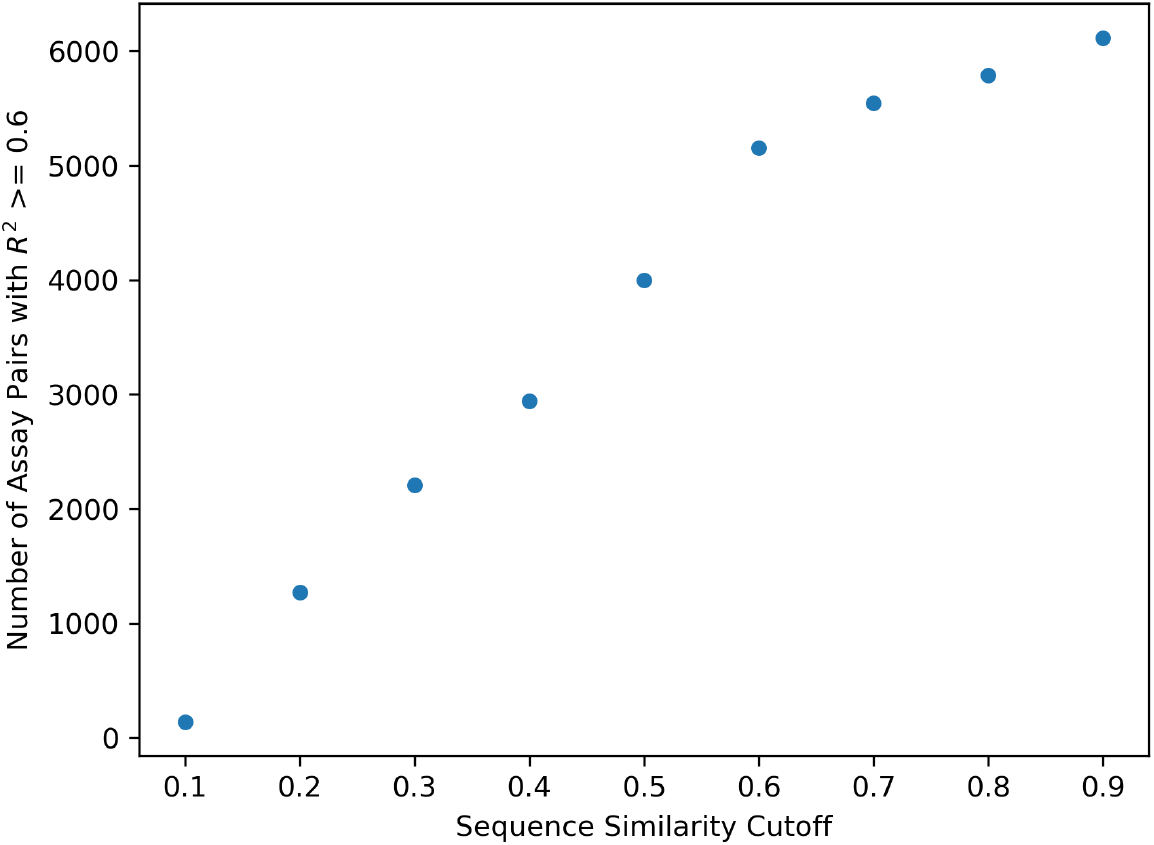
The relationship between sequence similarity and the number of assay pairs with correlated assays in the ChEMBL database. While the number of assay pairs drops as the similarity cutoff decreases, there are still more than 4,000 assay pairs remaining with a similarity cutoff of 0.5 (50%).

To ensure rigor in our train-test splits, we need to go beyond overall sequence similarity and consider both binding-site and ligand similarity. This approach was used by the team behind the PLINDER dataset [18] when creating their benchmark for co-folding performance. They accounted for overall sequence similarity, binding-site similarity, and ligand similarity when designing their benchmark dataset. Unfortunately, the task becomes more complex when designing a benchmark for activity prediction. Since we often lack information about where compounds bind, assessing binding-site similarity can be challenging or even impossible.

To assess model generalization, we propose separating the dataset into distinct ligand-similarity tiers. This directly controls for data leakage from ligand analogs, which has been shown to be a pervasive problem in the past [7] and in current benchmarks (see Sections 5 and 6). Inspired by the Runs N’ Poses benchmark [14] and based on ChEMBL 36 [22], we constructed the Novelty-Tiered Affinity Benchmark, which we make publicly available. The benchmark first splits the assays by publication year: assays from *≥* 2023 in test, 2022 in val, and *<* 2022 in train. We then create different compound novelty tiers by filtering each bin to contain only compounds within that Tanimoto similarity interval relative to the train and val sets. This is motivated by the fact that different use cases in drug discovery operate at different levels of compound novelty, from interpolation within a congeneric series (above 0.5 Tanimoto similarity) to the discovery of completely novel binders (below 0.35 Tanimoto similarity). Finally, we keep only assays with at least 10 compounds and a standard deviation of at least 0.5 across those compounds. See Appendix D for further details on this benchmark.

On this proposed benchmark, the best ligand-only performance drops considerably from *r* = 0.61 to *r* = 0.14 as we move from the least to the most stringent novelty tier (see Figure 4). This highlights a significant reduction in data leakage, particularly compared with the same model on the FEP+ 4 and OpenFE benchmarks (*r* = 0.66 and *r* = 0.36;

Figure 3). We also compared against a benchmark constructed using only a temporal split. Here, our best ligand-only method achieved *r* = 0.32, indicating that a temporal split alone is insufficient to prevent data leakage.

We filter the similarity bins by individual-compound similarity rather than by average assay-ligand similarity, as employed by IsoDDE [16] and Boltz-2 [15]. The challenge with average assay-ligand similarity is that it can allow partial leakage from high-similarity compounds into low-similarity assays. In Figure 8 in the Appendix, we show how the performance of the same ligand-only models varies when we bin by this method using the same bins as in the IsoDDE report. With this method, performance drops from *r* = 0.40 to *r* = 0.27 between the “high” and “medium” similarity bins, with the most stringent bin containing only 5 assays and exhibiting substantial variability.

The residual performance of the ligand-only models in Figure 4 likely stems from their learning general relationships between binding affinity and molecular properties within published series; for example, we found Pearson correlations of 0.19 and 0.082 between pIC_50_ and molecular weight and CLogP across the dataset.

## 8 Conclusions

The FEP+ 4 and OpenFE benchmarks, recently used to validate co-folding methods, contain severe instances of data leakage. The performance of a ligand-only baseline shows that ML models can achieve significantly inflated results by exploiting this leakage. Due to these limitations, this benchmark is not suitable for comparing different protein-ligand binding affinity methods. Claims that models like Boltz-2 and IsoDDE approach or exceed gold-standard physics-based methods were based on these benchmarks and should therefore be interpreted cautiously.

The problem is not specific to the 90% sequence-identity threshold used in these benchmarks. Our meta-analysis of 6,115 assay pairs in ChEMBL shows that even when overall sequence similarity is low, statistical correlations in assay activity remain high across homologous targets. As we showed by varying the similarity cutoff from 0.1 to 0.9, even a threshold as low as 0.5 leaves thousands of assay pairs for which activity can be predicted by cross-target mirroring. This systemic “target mirroring” in the medicinal chemistry literature renders overall protein-similarity methods, at any common threshold, unreliable for constructing binding-affinity benchmarks. Although clustering by binding-site similarity is an alternative, guessing the binding site for every interaction in the data set could introduce significant errors that invalidate the resulting comparisons.

This critique targets flawed benchmarks, not the models themselves. Large neural networks are known to memorize data, which does not preclude them from learning generalizable representations. However, these benchmarks provide no way to separate these contributions. Leakage-contaminated benchmarks make it impossible to separate memorization from genuine insight.

Machine learning methods for affinity prediction have repeatedly produced results that failed to transfer to prospective use. Building trust in these methods requires benchmarks that reflect the prospective task: predicting the affinity of genuinely novel compounds against targets of interest. To help the field in this direction, we propose and open-source the Novelty-Tiered Affinity Benchmark, which bins compounds by maximum Tanimoto similarity to the training or validation sets into high- and low-novelty tiers. We hope this will make it easier to separate models that rely on ligand memorization from those that learn generalizable representations. Finally, benchmarks shape model development; those with substantial data leakage create selection pressure favoring architectural choices that exploit it. We hope this contribution helps the field advance toward methods that truly enable chemists to discover novel binders.

## 9 Data and code availability

We make the Novelty-Tiered Affinity Benchmark publicly available at https://doi.org/10.5281/zenodo. 19665374 under CC BY-SA 3.0. The benchmark is built on ChEMBL 36. The code to evaluate models on this benchmark and produce the plot as in Figure 4 is made publicly available at https://github.com/bamattsson/ntab under the MIT license. This repository also contains code to regenerate the benchmark data and code for the ligand-only baseline methods. The code to reproduce figures in this paper is made available at https://github.com/bamattsson/paper-identifying_and_addressing_data_leakage under the MIT license.

## Appendix

### A FEP+ 4 and OpenFE test set reproducibility

To reproduce the FEP+ 4 and OpenFE test sets and data splits we followed these steps:

#### 1. Retrieve test set target seeds

- For FEP+ 4 the data points can be found in the github repo for protein-ligand-benchmark [28]. For the folders corresponding to each of the four targets TYK2, p38*α*, JNK1, CDK2 the corresponding 00_data/ligands.yml file was used to extract SMILES and experimental measurements.
- For the OpenFE dataset data points have been published in an accessible form by the Open Free Energy initiative [29] (this is the dataset cited by IsoDDE and contains 873 measurements, Boltz-2 uses a slightly different dataset with 867 measurements). Mapping the `system group` and `system name` columns to targets can be achieved via the `industry_benchmarks/input_structures/original_structures/*/subset_metadata.csv` files in the same repository and the `Reference PDB` entries.

#### 2. Find protein sequences

Protein sequences were downloaded from ChEMBL (version 36) [22]; two test targets not present in ChEMBL were retrieved from UniProt [30].

#### 3. Clustering

We clustered protein sequences using MMseqs2 [31] following the protocol in [15]: `mmseqs easy-cluster fasta_seqs_for_clustering.fasta cluster_res cluster_tmp --min-seq-id 0.9 --cov-mode 0 -c 0.01`.

#### 4. Splitting the data

Protein targets sharing a cluster with any FEP+ 4 or OpenFE test target were assigned to the test set. The remaining clusters were allocated to the training or validation sets.

### B Further results on the FEP+ OpenFE benchmarks

#### Data leakage in the FEP+ OpenFE subset

Figure 6 shows additional data leakage problems in the FEP+ OpenFE subset when using a 90% sequence-identity threshold to split the dataset into train and test targets. For both of these cases, the target in the test set has a close analogue in the training set, which has sequence identity below the threshold.

**Figure 6:**
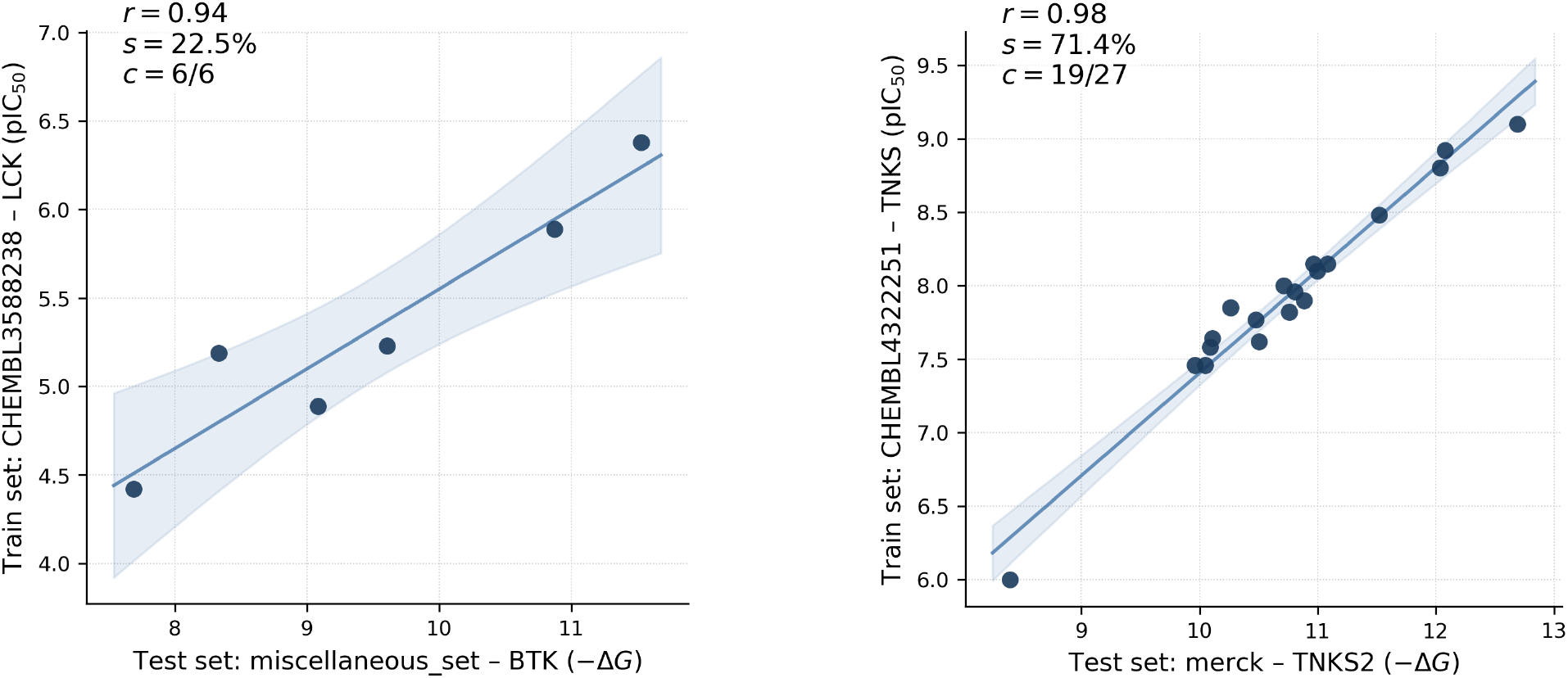
Data leakage in the FEP+ OpenFE subset when using a 90% sequence identity. The x-axes contain the test data points with experimental *™* Δ*G*, and y-axes show pIC_50_ measurements of the same compounds that appear in the training split for closely related targets. *r* shows the Pearson correlations, *s* the sequence identity between the two targets, and *c* the coverage of compounds from the test assay that also appear in the training assay.

#### Reproducing the Boltz-2 compound similarity analysis

The Boltz-2 paper analyzes how affinity prediction performance on the FEP+ OpenFE test set depends on the similarity of compounds to their training sets [15]. The idea is that failing to show a relationship between compound similarity and performance would indicate that the model is not reliant on similar compounds in the training data. To assess whether this test has sufficient statistical power, we reproduced it with the chemprop ligand-only baseline. We know that ligand-only models rely on interpolating between seen ligand structures; if the test is powerful enough, we should be able to detect this. Figure 7 shows the results. A Tukey-HSD test failed to reject the null hypothesis that there is no difference among the bins. Hence, this analysis is not powerful enough to identify methods that work through ligand memorization. One reason for this could be that there are not many data points in each bin: 10–21 compared with 24–340 in the benchmark we propose. Another reason could be that some of these have very few data points; the smallest contains only 3, leading to larger noise levels within each bin. We should note that this dataset is not exactly the same as the one used in the Boltz-2 report; they report using 867 measurements, whereas we have used 873 (See Appendix A).

**Figure 7:**
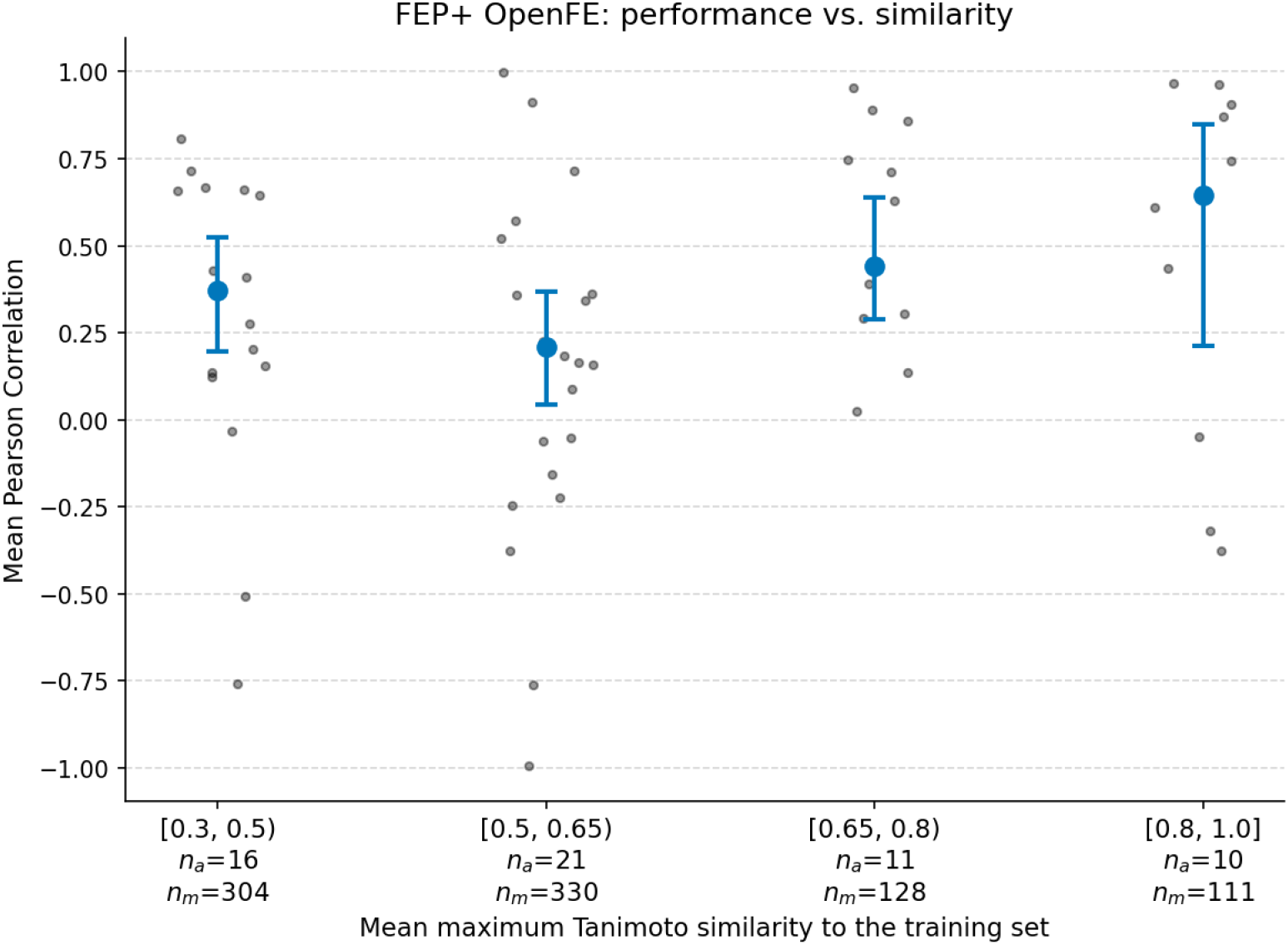
Relationship between compound similarity and prediction performance on the OpenFE benchmark for the ligand-only chemprop model. Each assay is assigned to a bucket depending on the average maximum Tanimoto similarity to all training compounds. *n*_*a*_ is the number of assays and *n*_*m*_ number of measurements in that bucket. Error bars reflect bootstrapped 95% confidence intervals. Tukey-HSD fails to reject the null hypothesis that there is no difference between any of the buckets.

### C Ligand-only baseline model

The baselines are ligand-only models trained to predict pIC_50_, p*K*_i_ and p*K*_d_ values from ligand features and a learned target representation, with no structural information about the protein target. This model represents a lower bound on what is achievable via ligand interpolation alone; more advanced ligand-only models with better representations of the ligand pharmacophore could achieve higher performance.

#### Inputs

Each training example consists of up to four inputs: a molecular graph (atom and bond featurized using Chemprop [26]), a Morgan Fingerprint (2048-bit, radius 2, log-transformed counts), a vector of *N*_prop_ normalised molecular descriptors (Table 4), and an integer target index.

**Table 4:** Molecular descriptors used as input features to the baseline model, computed with RDKit.

|  |  |  |  |
| --- | --- | --- | --- |
| Lipophilicity (CLogP) | Mol. weight (Da) | TPSA ($\text{\AA}^2$) | H-bond donors |
| H-bond acceptors | Rotatable bonds | Formal charge | Molar refractivity |
| Fraction sp<sup>3</sup> C | Ring count | Aromatic ring count | Heavy atom count |

#### Architecture

The molecular graph is encoded by a Chemprop Directed Message Passing Neural Network (D-MPNN) consisting of a bond message passing network with hidden dimension *d* and depth *n*_*h*_, followed by a Chemprop NormAggregation to produce a molecule-level vector. Optionally, a Morgan Fingerprint is projected through a single linear layer with batch normalisation (BN) and Gaussian Error Linear Unit (GELU) activation to produce a *d*-dimensional encoding. The available encodings are concatenated with the molecular descriptors vector (if enabled) and a learned target embedding of dimension *d*_target_, yielding a combined representation that is fed to a two-layer Multi-Layer Perceptron (MLP) head. The final scalar prediction adds two learned bias terms: one per target and one per assay type (pIC_50_, p*K*_i_ or p*K*_d_). The full architecture is summarised in Table 5.

**Table 5:**
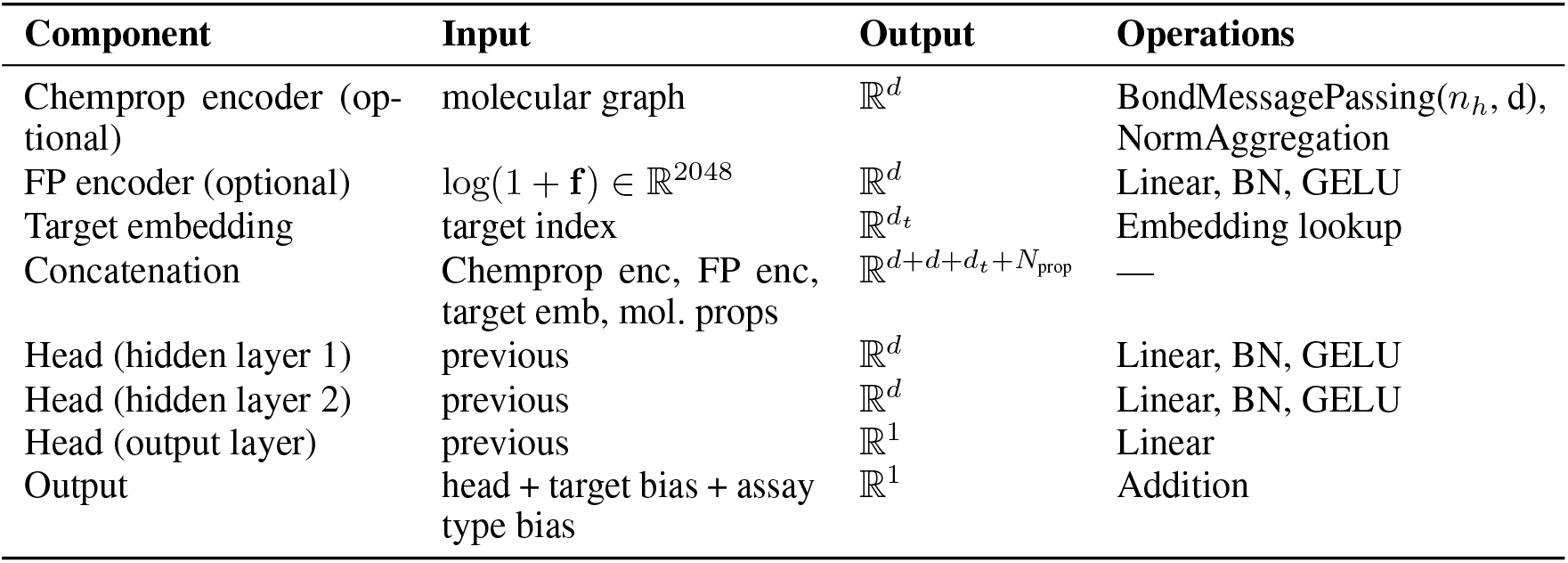
Architecture of the ligand-only baseline models. *d* = *d*_hidden_ = 2048, *d*_*t*_ = *d*_target_ = 256, *n*_*h*_ = *n*_hidden_ = 4. **F** is the Morgan Fingerprint counts (2048-bit, radius 2).

#### Target embedding

The target embedding is an integer lookup with no structural input, allowing the model to learn which compounds bind to which targets from historical data, but, by design, preventing it from generalizing to genuinely novel binding environments (such as unexplored regions of existing pockets, new pockets, or compounds without analogs). For targets absent from the training set, the embedding and bias of the nearest training target by protein sequence similarity were used.

#### Training

The model was trained with Adam using a Noam-like learning rate schedule: the learning rate increases linearly from 10^*−*4^ to 10^*−*3^ over the first two warmup epochs, then decreases linearly to 10^*−*4^ over the remaining epochs. Training ran for 100 epochs with the best checkpoint selected by validation Pearson *r*.

### D Benchmark construction method

The benchmark is constructed from ChEMBL 36 [22]. We retain only high-confidence binding assays against single protein targets, restricting to maximum confidence score (9), binding assay type, direct or homologous target mapping, and the three standard affinity readouts (*K*_i_, *K*_d_, IC_50_). Flagged or unreliable measurements and assays against mutant sequences are excluded. All affinity values are converted to the *−* log_10_ scale. Activities with both equality- and inequality-relation measurements are kept in the training set. Activities flagged as potential duplicates are also kept as some of this data is genuinely novel. The user can choose to filter these out if they wish to.

#### Time split

Assays are partitioned by the publication year of their source document in ChEMBL. Assays published before 2022 form the training set; those published in 2022 the validation set; and those published from 2023 onward are candidates for the test set. Since the validation set influences model development through early stopping and hyperparameter optimization, leakage across the test/val boundary would inflate reported performance in the same way as leakage across test/train. By using a cutoff date for the test set, model builders who choose a two-step approach of both co-folding and binding prediction can minimize risk of data leakage in either of the two steps by aligning cutoff dates for the two models.

#### Novelty-tiers

Time-split on its own is insufficient, as most assays published in a given year include compounds that have appeared in earlier literature. To control for this leakage we compute the maximum Tanimoto similarity (2048-bit Morgan fingerprints with radius 2) between each new compound and all compounds whose earliest ChEMBL publication predates the cutoff date. From this we construct different novelty-tiers of compounds with a certain maximum similarity, and calculate the performance within those tiers. We also experimented with binning by average maximum Tanimoto similarity per assay, rather than by individual compound similarity. Figure 8 shows the results. The difference is less stark than in Figure 4, reflecting the fact that this method yields low-similarity assays that still might contain high-similarity compounds. We should note that even though we have here attempted to reproduce the plot shown in the IsoDDE report there are several differences: they use ChEMBL 35, they use data from 2023 and onwards both for val and test, they use more activity types, they filter to assays with at least 20 ligands.

**Figure 8:**
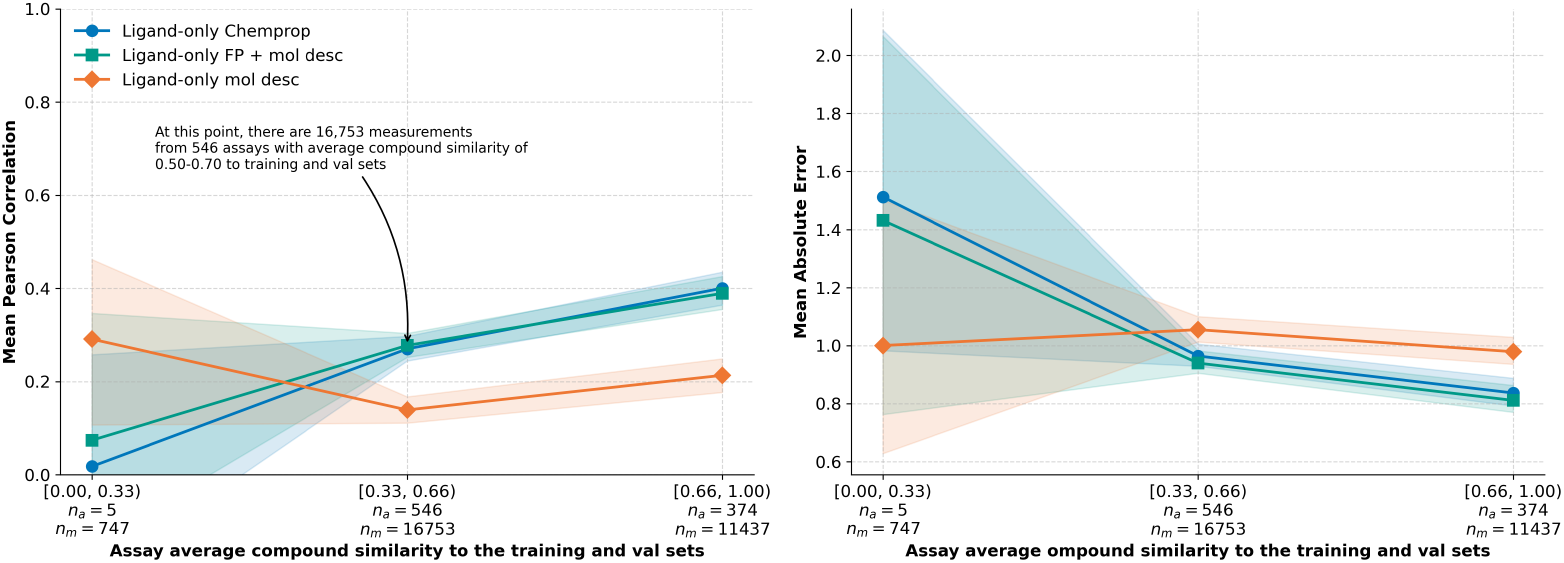
Performance of ligand-only baseline models for mean maximum ligand novelty tiers. Each bin along the x-axis groups assays by their average maximum compound Morgan Fingerprint (2048-bit, radius 2) Tanimoto similarity to training or validation compounds. *n*_*a*_ is the number of assays and *n*_*m*_ the number of measurements within that bin. The metric mean was calculated as an unweighted average across all assays in that bin, and shaded areas show bootstrapped 95% confidence intervals. For all three models the changes between the [0.33, 0.66) and [0.66, 1.00) bins were statistically significant with Tukey-HSD, for the Chemprop model the difference between [0.00, 0.33) and [0.66, 1.00) were also statistically significant.

#### Assay predictability filter

We retain from the test and validation sets only assays with at least 10 unique compounds, a *−* log_10_ standard deviation of at least 0.5 across those compounds, and equality-relation measurements only. Correlation coefficients against assays with insufficient compounds or low-affinity variations would be dominated by measurement noise. Finally, to avoid biasing the benchmark with multiple near-identical series from the same experimental campaign, we retain at most one assay per document, keeping the largest qualifying assay when a document contributes multiple.

